# Heparan sulfate assists SARS-CoV-2 in cell entry and can be targeted by approved drugs *in vitro*

**DOI:** 10.1101/2020.07.14.202549

**Authors:** Qi Zhang, Catherine Z. Chen, Manju Swaroop, Miao Xu, Lihui Wang, Juhyung Lee, Amy Q. Wang, Manisha Pradhan, Natalie Hagen, Lu Chen, Min Shen, Zhiji Luo, Xin Xu, Yue Xu, Wenwei Huang, Wei Zheng, Yihong Ye

## Abstract

The cell entry of SARS-CoV-2 has emerged as an attractive drug repurposing target for COVID-19. Here we combine genetics and chemical perturbation to demonstrate that ACE2-mediated entry of SARS-CoV and CoV-2 requires the cell surface heparan sulfate (HS) as an assisting cofactor: ablation of genes involved in HS biosynthesis or incubating cells with a HS mimetic both inhibit Spike-mediated viral entry. We show that heparin/HS binds to Spike directly, facilitates the attachment of viral particles to the cell surface to promote cell entry. We screened approved drugs and identified two classes of inhibitors that act via distinct mechanisms to target this entry pathway. Among the drugs characterized, Mitoxantrone is a potent HS inhibitor, while Sunitinib and BNTX disrupt the actin network to indirectly abrogate HS-assisted viral entry. We further show that drugs of the two classes can be combined to generate a synergized activity against SARS-CoV-2-induced cytopathic effect. Altogether, our study establishes HS as an attachment factor that assists SARS coronavirus cell entry, and reveals drugs capable of targeting this important step in the viral life cycle.

---

The ongoing pandemic of the coronavirus disease 2019 (COVID-19) has claimed many lives and severely damaged the global economy. This severe acute respiratory syndrome (SARS) is caused by a novel coronavirus SARS-CoV-2 ^1, 2^, which is closely related to SARS-CoV, the virus underlying the 2003 SARS outbreak ^3^.

As a positive-sense, single-stranded RNA virus bearing a membrane envelope, coronavirus enters cells when this membrane envelope fuses with host membranes ^4^. Viral entry may take place either at the plasma membrane or at the endosomes following receptor-mediated endocytosis ^5^. Previous studies showed that SARS-CoV enters cells primarily through ACE2-dependent endocytosis ^5-9^. New evidence suggests that SARS-CoV-2 may follow the same entry path ^1, 10, 11^. Specifically, limiting the production of phosphatidylinositol 4,5-bisphosphate (PIP2) inhibits the fusion of the SARS-CoV-2 envelop with the endolysosomes and viral entry ^12, 13^. Additionally, viral entry-associated membrane fusion requires a priming step mediated by several host proteases including the lysosome-localized Cathepsin B/L and serine proteases of the TMPRSS family ^12, 14-18^. Although some TMPRSS proteases may act on the cell surface, the fact that increasing the lysosomal pH in either TMPRSS2 positive Caco-2 or TMPRSS2 negative HEK293 cells both inhibits SARS-CoV-2 cell entry ^14^ suggests endolysosomes as a major entry site for SARS-CoV-2 at least in certain cell types.

Our previous study on the intercellular transmission of misfolded α-Synuclein (α-Syn) fibrils, a cellular process reminiscent of viral infection, revealed an endocytosis mechanism by which the cell surface heparan sulfate (HS) facilitates receptor-mediated uptake of protein assemblies bearing excess positive charges ^19^. HS is a negative charge-enriched linear polysaccharide molecule that is attached to several membrane and extracellular proteins, which are collectively termed as heparan sulfate proteoglycans (HSPG). The cell surface HS can serve as an anchor point to facilitate endocytosis of many cargos ^20, 21^, which include SARS-CoV-2 related coronaviruses ^22-25^. To identify drugs that can diminish the spreading of misfolded α-Syn and the associated Parkinsonism ^26^, we screen and identify eight FDA-approved drugs that block the HS-dependent endocytosis of preformed α-Syn fibrils (see below). Intriguingly, one of the candidates Tilorone was recently reported as an inhibitor of SARS-CoV-2 infection ^27^. This coincidence, together with the recently reported interaction of SARS-CoV-2 Spike with the HS mimetic glycan heparin ^28-30^ and other evidence in support of a role for HS in the entry of SARS-CoV-2-related coronaviruses ^22-25, 31^ prompted us to investigate the possibility of targeting HS as a COVID-19 therapeutic strategy.

## Results

### Heparan sulfate facilitates Spike-dependent viral entry

To study Spike-mediated viral entry, we constructed luciferase-expressing pseudoviral particles (PP) bearing SARS-CoV or SARS-CoV-2 Spike. We infected HEK293T (human embryonic kidney cells) cells or HEK293T cells stably expressing ACE2-GFP with PP. As expected, ACE2-GFP HEK293T cells transduced with SARS-CoV or SARS-CoV-2 PP expressed luciferase significantly higher than non-infected ACE2-GFP cells (Fig. S1A, B) or infected HEK293T cells without ACE2-GFP (Fig. S1C). These results confirm Spike-dependent entry of SARS-CoV and CoV-2 via ACE2.

To test the role of HS in viral entry, we first treated ACE2-GFP HEK293T cells with heparin, a HS mimetic glycan frequently used as a competitive inhibitor for HS ligands (Figure 1A) ^32, 33^. Heparin treatment dose-dependently mitigated luciferase expression from both SARS-CoV and CoV-2 PP (Figure 1B, C) with little impact on cell viability (Fig. S1D). Heparin treatment also reduced luciferase expression in SARS-CoV- or CoV-2-transduced human lung epithelial Calu-3 cells (Fig. S1E) and the level of inhibition was more significant than in ACE2-GFP overexpressing cells. Heparin pulldown showed that the purified Spike ectodomain readily bound to heparin-conjugated beads (Figure 1D). As expected, the interaction was sensitive to salt, but a significant amount of Spike remained associated with heparin in a buffer containing 120mM NaCl (Fig. S1F), a condition recapitulating the airway surface salt concentration ^34^. These data suggest that heparin acts as a competitive inhibitor for Spike-mediated viral entry and this inhibitory activity could be offset by overexpression of ACE2.

**Figure 1.**
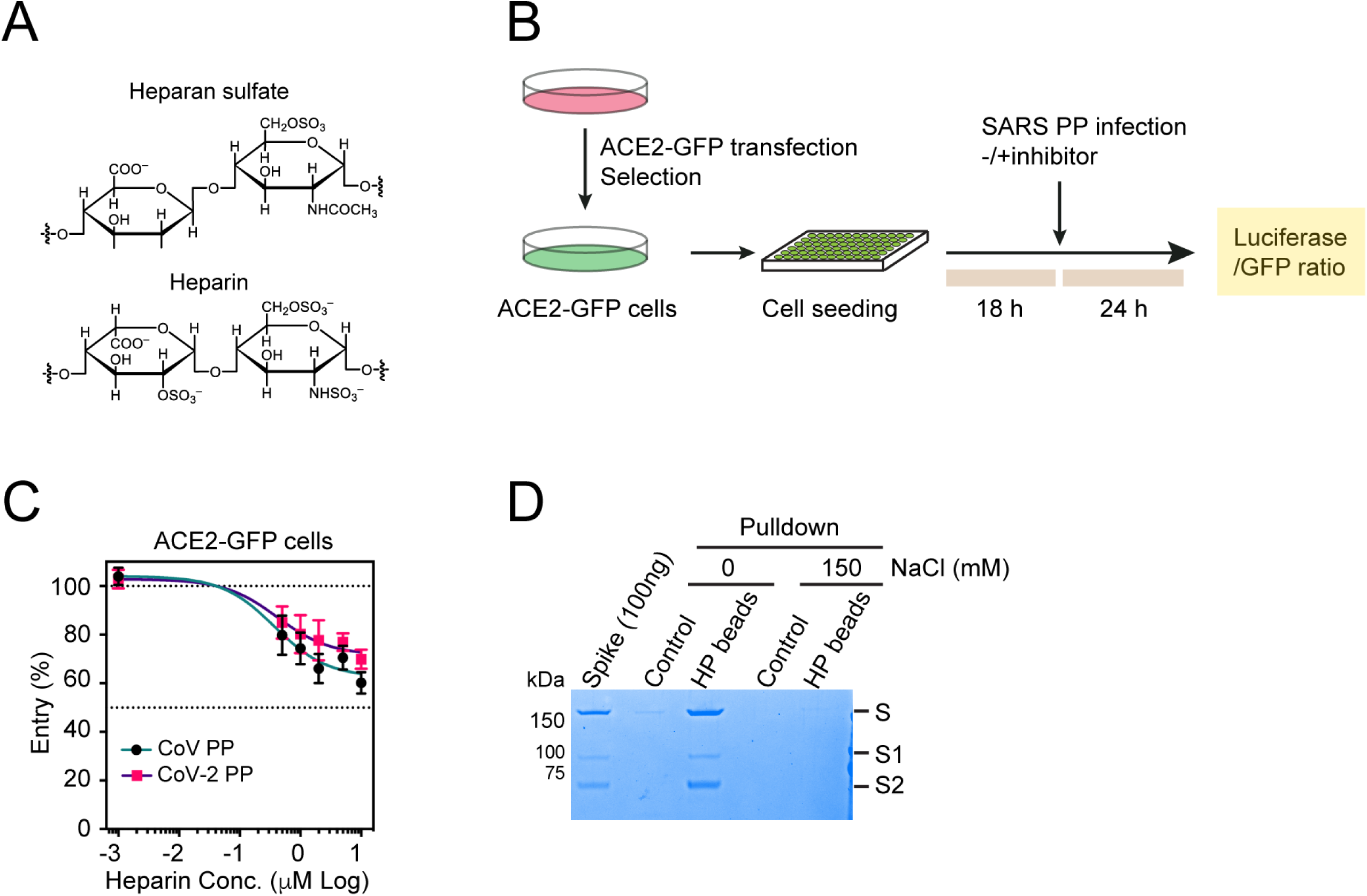
Heparin inhibits Spike-mediated SARS-CoV and CoV-2 entry. **(A)** The chemical structures of heparan sulfate and heparin. **(B)** The experimental scheme for inhibitor testing in HEK293T ACE2-GFP cells. **(C)** Heparin mitigates the entry of SARS-CoV and SARS-CoV-2 pseudoviral particles (PP). ACE2-GFP HEK293T cells were transduced with SARS-CoV and SARS-CoV-2 PP in the presence of heparin as indicated. The ratio of luciferase vs. GFP was measured 24 h post-transduction. Error bars indicate SEM, N=4. **(D)** Heparin interacts with Spike in a salt sensitive manner. Spike (600ng) was incubated with either control or heparin-conjugated beads in a buffer containing 0 or 150mM NaCl. Bound proteins were analyzed by SDS-PAGE and Coomassie blue staining. Note that a small amount of Spike interacts with heparin in the presence of 150mM NaCl.

Next, we used small interfering RNA (siRNA) or CRISPR to disrupt genes encoding HSPG biosynthetic enzyme XYLT2 and SLC35B2 ^19^ (Figure 2A, Fig. S1G-I). *XYLT2* encodes one of the two HS chain initiation enzymes. SLC35B2 is a Golgi-localized transporter for 3’-phosphoadenosine 5’-phosphosulfate (PAPS), which is essential for HS chain sulfation ^32^. Knockdown of *XLYT2* by ∼80% inhibited SARS-CoV and SARS-CoV-2 PP entry similarly as heparin treatment (Figure 2B). By contrast, CRISPR-mediated inactivation of *SLC35B2* completely abolished HSPG biosynthesis ^19^ and thus inhibited the entry of SARS-CoV and SARS-CoV-2 more dramatically (Figure 2C). Analyses of GFP fluorescence showed no effect of *SCL35B2* knockout on ACE2-GFP expression (Fig. S1J). Nevertheless, the knockout of *SLC35B2* significantly reduced the binding of SARS-CoV-2 PP to ACE2-GFP cells (Figure 2D, E). Altogether, these results support a model in which the cell surface HS serves as a virus-recruiting factor to promote ACE2-dependent viral entry.

**Figure 2.**
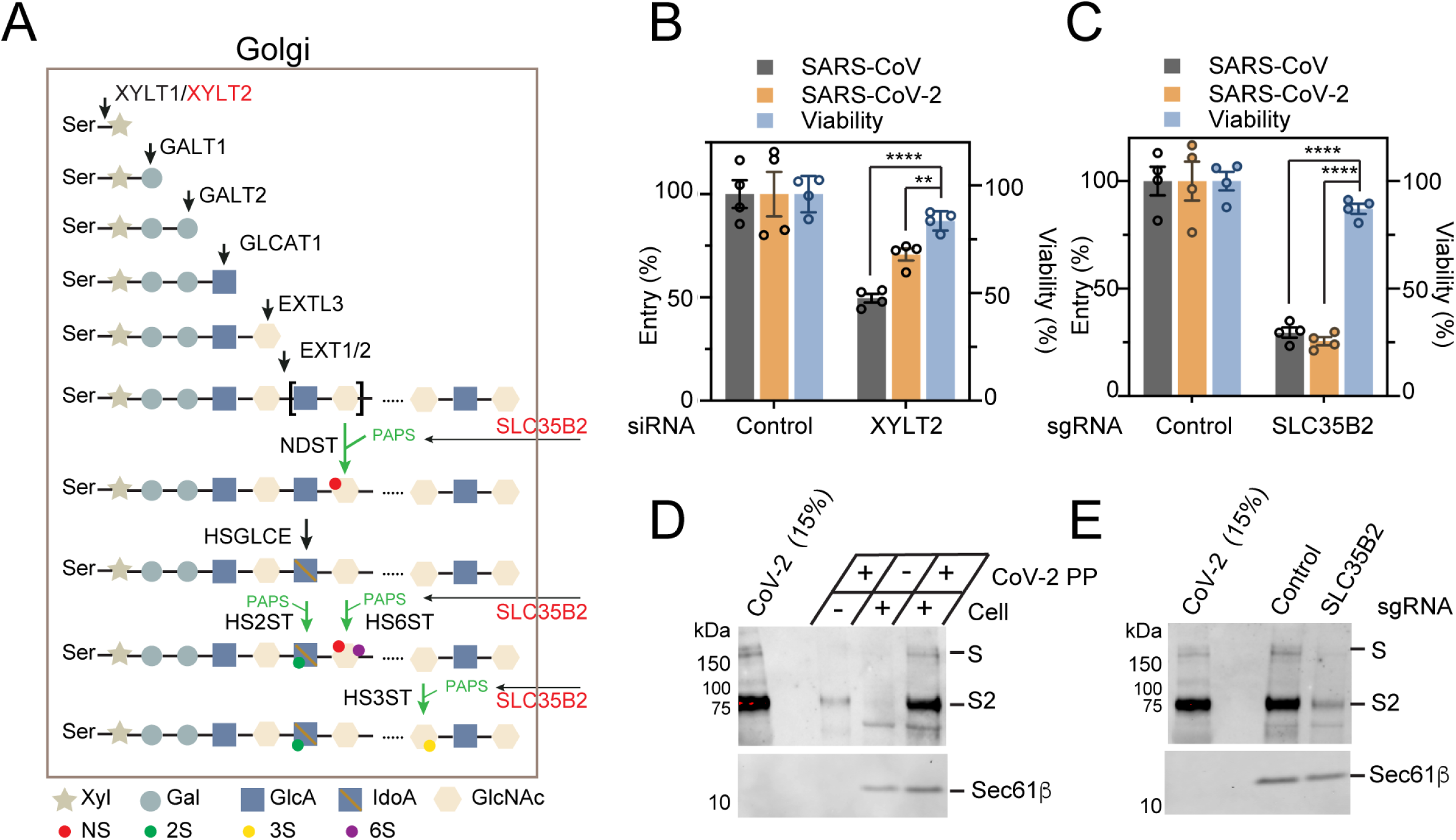
Heparan sulfate promotes Spike-mediated SARS-CoV and CoV-2 entry. **(A)** The HSPG biosynthetic pathway. Genes chosen for knockdown or knockout (KO) are in red. **(B)** Knockdown of *XYLT2* reduces SARS-CoV and SARS-CoV-2 PP entry. ACE2-GFP cells transfected with either control or *XYLT2* siRNA were transduced with SARS-CoV (grey) or SARS-CoV-2 (orange) PP for 24 h and the ratio of luciferase/GFP was determined. A parallel experiment done without the virus provides another control for the effect of gene knockdown on cell viability (blue). Error bars indicate SEM, N=4. **, *p*<0.01, ****, *p*<0.0001 by unpaired Student t-test. **(C)** *SCL35B2* is required for SARS-CoV and SARS-CoV-2 cell entry. As in B, except that control and *SLC35B2* CRISPR KO cells were used. (**D** and **E**) *SLC35B2* promotes the binding of SARS-CoV-2 PP to cells. (D) ACE2-GFP cells were spin-infected at 4 °C for 1 h. After washing, the virus bound to the cells was detected by immunoblotting. (E) The binding of SARS-CoV-2 PP to control and *SLC35B2* deficient cells was analyzed by immunoblotting. S and S2 indicate the full length and furin-cleaved S2 fragment, respectively.

### A drug screen identifies inhibitors targeting the HS-dependent cell entry pathway

HS is a negatively charged biopolymer, which can recruit ligands bearing positive charges to facilitate their endocytosis ^20^. We recently characterized the endocytosis mechanism of α-Syn fibril, which is a HS ligand ^19^. We then conducted a quantitative high-throughput (qHTS) screen using α-Syn filamentous inclusions to search for approved drugs that could block HS-dependent endocytosis (Figure 3A). The screen identified 8 drugs that inhibited α-Syn fibril uptake in HEK293T cells (Figure 3B). Additional studies using a panel of endocytosis cargos confirmed that the identified drugs are endocytosis inhibitors with a preference for HS-dependent ligands (Figure 3C). Among the cargos tested, supercharged GFP (GFP+), polycation-coated DNA, and VSVG-pseudotyped lentivirus all enter cells via a *SLC35B2* dependent mechanism ^19, 35^, whereas Transferrin uses a HS independent but clathrin-dependent endocytosis pathway. The cholera toxin B chain (CTB) reaches the Golgi apparatus via a clathrin-independent mechanism ^36^. None of the drugs tested affected the internalization of CTB, but Sunitinib and BNTX appeared to disrupt the perinuclear stacking of the Golgi system (Fig. S2A, B). These drugs also had little impact on Transferrin endocytosis (Fig. S2C), but they could reduce the uptake of HS-dependent cargos to various levels with Mitoxantrone being the most potent one: at 5 μM, it almost completely blocked the endocytosis of all HS-dependent cargos tested (Figure 3C, Fig. S2D-F).

**Figure 3.**
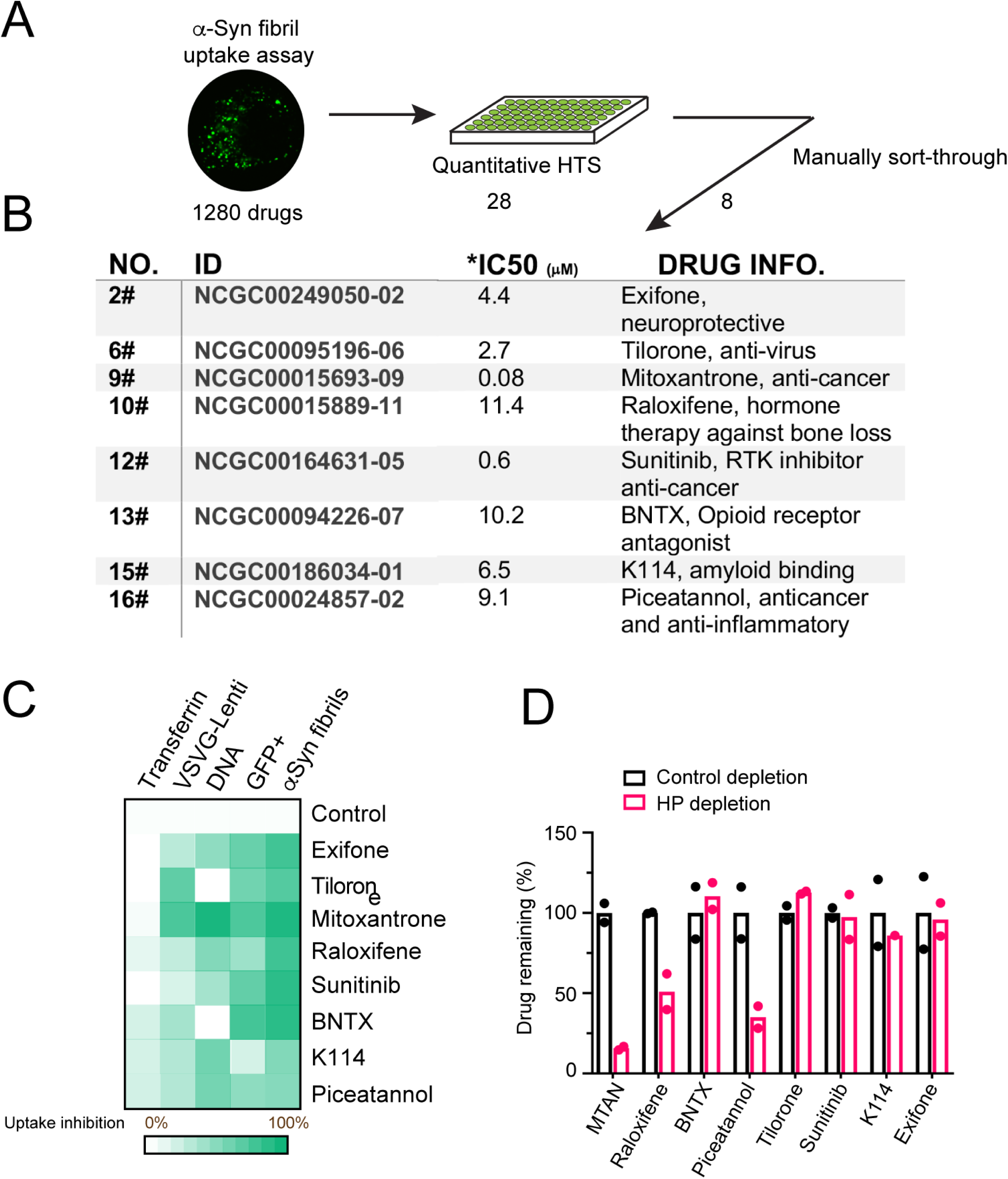
Two classes of drugs targeting HS-dependent cargo entry. **(A)** A high-throughput drug screen for inhibitors that block the uptake of fluorescently labeled α-Syn fibrils. **(B)** A summary of the drugs identified. AC_50_ was calculated from the automated screen. IC_50_ was determined by a follow-up repeat shown in D. **(C)** A heat map summary of the endocytosis inhibitory activity for the indicated drugs. All drugs were tested at 10 μM except for Sunitinib and Mitoxantrone (5 μM each). See Supplementary Figure 2 for details. **(D)** Mass spectrometry analysis of drug binding to heparin beads. The indicated drugs were incubated with either control or heparin-conjugated beads. Unbound drugs were quantified by mass spectrometry. N=2. MTAN, Mitoxantrone.

Biochemical binding coupled to mass spectrometry analyses showed that these drugs could be categorized into two classes depending on whether they bound to heparin. While Exifone, K114, Tilorone, BNTX, and Sunitinib showed no affinity to heparin, Mitoxantrone, Raloxifene, and Piceatannol in solution could be effectively depleted by heparin beads (Figure 3D), suggesting that these drugs amy target the cell surface HS directly to inhibit cargo uptake.

Among the drugs identified, Tilorone was previously established as a pan-antiviral agent ^37^ that also inhibits SARS-CoV-2 infection *in vitro* ^27^. This coincidence, together with the convergence of SARS-CoV and CoV-2 entry and α-Syn endocytosis at the HS glycans raise the possibility that inhibitors of HS-dependent endocytosis might also block SARS-CoV and CoV-2 entry. Indeed, with the exception of Exifone and K114, other drugs could all mitigate Spike-mediated viral entry in ACE2-GFP cells (Fig. S3A and see below). Exifone and K114 were excluded from further study.

### The actin network is required for Spike-mediated viral entry

We chose representative drugs of the two classes for further characterization. For those that did not bind heparin, we characterized BNTX and Sunitinib because their effect on Golgi morphology suggested a potential shared mechanism. BNTX is an opioid receptor antagonist for opioid or alcohol use disorders, while Sunitinib has been used as a receptor tyrosine kinase inhibitor for cancer therapy ^38^. In HEK293 ACE2-GFP cells treated with SARS-CoV or SARS-CoV-2 PP, these drugs inhibited viral entry with respective IC_50_ of ∼30μM and 10μM for Sunitinib, 9μM and 10μM for BNTX (Figure 4A, B). In Vero E6 cells treated with wild-type SARS-CoV-2, BNTX at 2 μM rescued the virus-induced CPE by ∼70%, whereas 10μM Sunitinib rescued the CPE only by ∼30% (Figure 4C, Fig. S3B).

**Figure 4.**
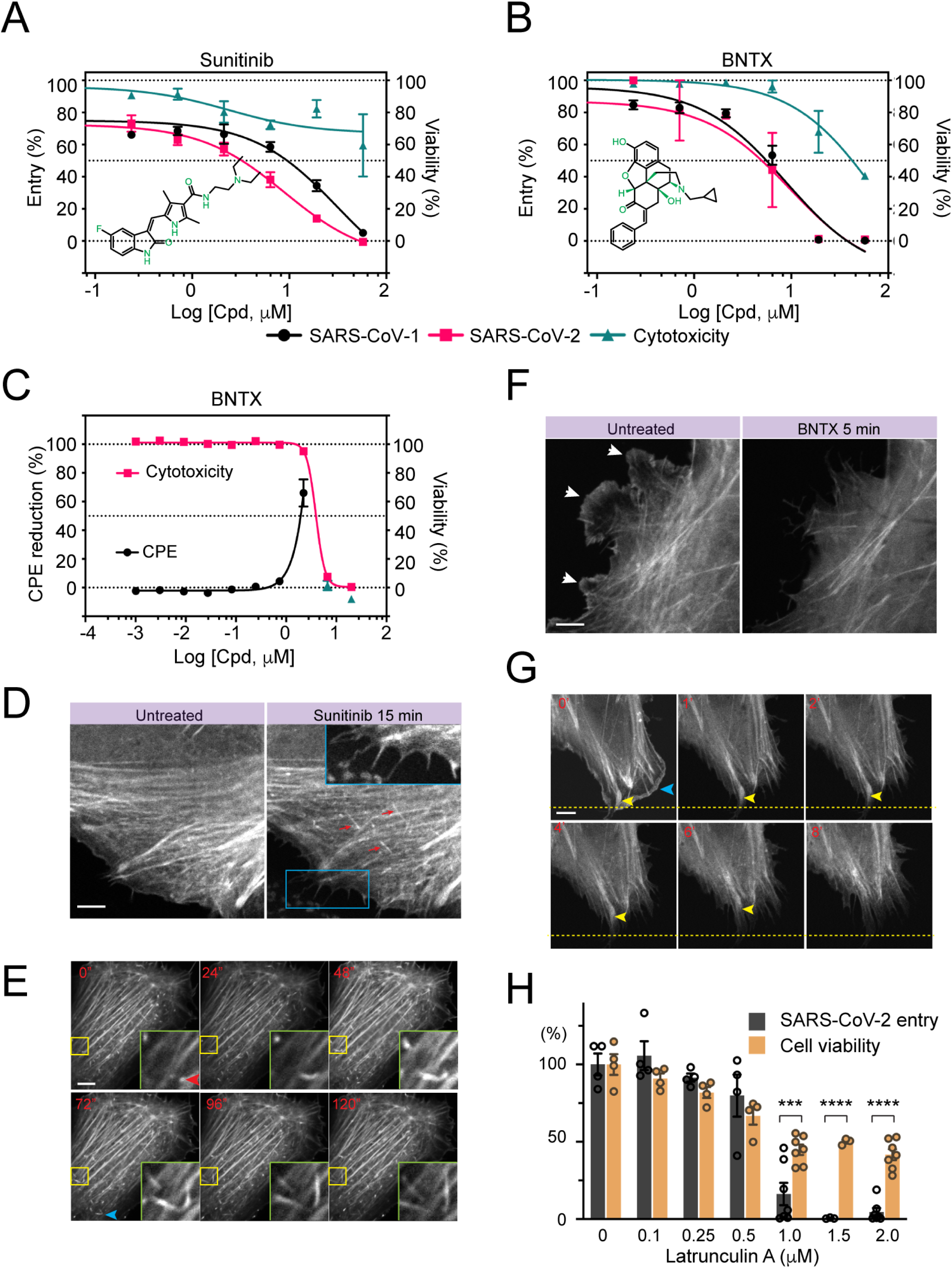
The actin cytoskeleton is required for HS-assisted viral entry. (**A, B**) Sunitinib and BNTX inhibit Spike-dependent entry of pseudoviral particles (PP). HEK293 ACE2-GFP cells were transduced with SARS-CoV and SARS-CoV-2 PP in the presence the indicated drugs. The luciferase expression was measured 48 h post-transduction. Error bars indicate SEM, N=4. As a control for cytotoxicity, cells treated with the drugs without virus were analyzed in parallel by an ATP-based cytotoxicity assay. (**C**) BNTX protects Vero E6 cells from SARS-CoV-2-induced cytopathic effect (CPE). Viability of Vero E6 cells was measured after treatment with the indicated drugs in the presence (black) or absence (red) of the SARS-CoV-2 virus. Error bars indicate SEM, N=2. (**D** and **E**) Sunitinib stimulates actin filament formation at the cell periphery. (D) Confocal snapshots of a U2OS cell stably expressing Tractin-EGFP before or after Sunitinib (5 μM) treatment. The red arrows indicate new actin filaments formed in a direction perpendicular to the existing filaments. The inset shows an enlarged view of the box, which highlights newly formed filopodia. (E) Live cell imaging of actin filament formation after Sunitinib treatment (5 μM, 60min). The arrows indicate examples of actin filament assembly. The insets show an enlarged view of the boxed area. Scale bar, 5 μm. (**F** and **G**) BNTX disrupts actin filaments at the cell periphery. (F) Confocal snapshots of a Tractin-EGFP cell before and after BNTX (10 μM) treatment. Arrows indicate the peripheral actin network disrupted by BNTX. (G) Live cell imaging shows the shrinking of peripheral membranes after BNTX treatment. Yellow arrowheads mark a retracting actin bundle. (**H**) Latrunculin A inhibits SARS-CoV-2 entry. ACE2-GFP cells incubated with SARS-CoV-2 PP in the presence of Latrunculin A were analyzed for luciferase expression (Entry). A parallel treatment in the absence of the virus showed the toxicity of Latrunculin A (orange). Error bars indicate SEM. N=2. ***, *p*<0.001, ****, *p*<0.0001.

Because HS-assisted endocytosis often requires the actin cytoskeleton ^19^, we determined whether these drugs affect the actin cytoskeleton network using U2OS cells stably expressing Tractin-EGFP, an actin filament label ^39^. Confocal microscopy showed that actin monomers polymerize near peripheral membranes in untreated cells, forming a microfilament network with parallel actin filaments (Fig. S3C). Additionally, a meshwork of actin is associated with the cell cortex, while thick actin bundles named stress fibers are often found near basal membranes. In cells exposed to Sunitinib, the actin microfilaments were frequently replaced by short disoriented actin segments. Many cells also contained an increased number of filopodia. By contrast, cells treated with BNTX had significantly reduced number of actin filaments, with many cells containing actin-positive aggregates (Fig. S3C, top panels). These phenotypes were further confirmed by TIRF microscopy in Sunitinib- or BNT-treated cells but were not observed in cells treated with other drugs (Fig. S3C, D).

To further elucidate the actions of Sunitinib and BNTX, we used live-cell imaging to monitor the acute changes in actin dynamics in drug-treated cells. Shortly after Sunitinib treatment, we observed extensive de novo actin filament formation (Figure 4D, E, Supplementary movies 2 vs. 1). Actin polymerization was also detected in the cell periphery where it drove filopodium formation (Figure 4D) but was not seen near basal membranes where stress fibers were located (Supplementary movie 4 vs. 3). Unlike Sunitinib, BNTX treatment resulted in rapid shrinking of actin microfilaments and the loss of membrane ruffling in the peripheral cortex (Figure 4F, G, Supplementary movies 6 vs. 5). These data suggest that Sunitinib promotes actin assembly to form disoriented filaments, whereas BNTX disrupts the cortex actin meshwork either by promoting actin disassembly or inhibiting actin assembly. The fact that two structurally unrelated actin inhibitors both block SARS-CoV and CoV-2 entry strongly suggests that the entry of these viruses may require the actin cytoskeleton. Accordingly, SARS-CoV-2 PP entry was also inhibited by Latrunculin A, a well-established actin inhibitor (Figure 4H). These findings, together with the fact that Sunitinib was previously identified as an entry inhibitor for Ebola virus ^40^ highlight an important role played by actin in the entry of these RNA envelop viruses.

### Mitoxantrone inhibits viral entry by targeting HS

Among drugs bound to heparin, Mitoxantrone had a higher affinity. Mitoxantrone was previously reported as a DNA intercalator, which also inhibits type II DNA topoisomerase ^41, 42^. Accordingly, Mitoxantrone is currently approved for treatment of acute non-lymphocytic leukemia, prostate cancer, and multiple sclerosis. In PP-treated ACE2-GFP HEK293 cells, Mitoxantrone strongly inhibited viral entry with IC_50_ >100-fold lower than that of cytotoxicity, suggesting an excellent anti-viral therapeutic window (Figure 5A).

**Figure 5.**
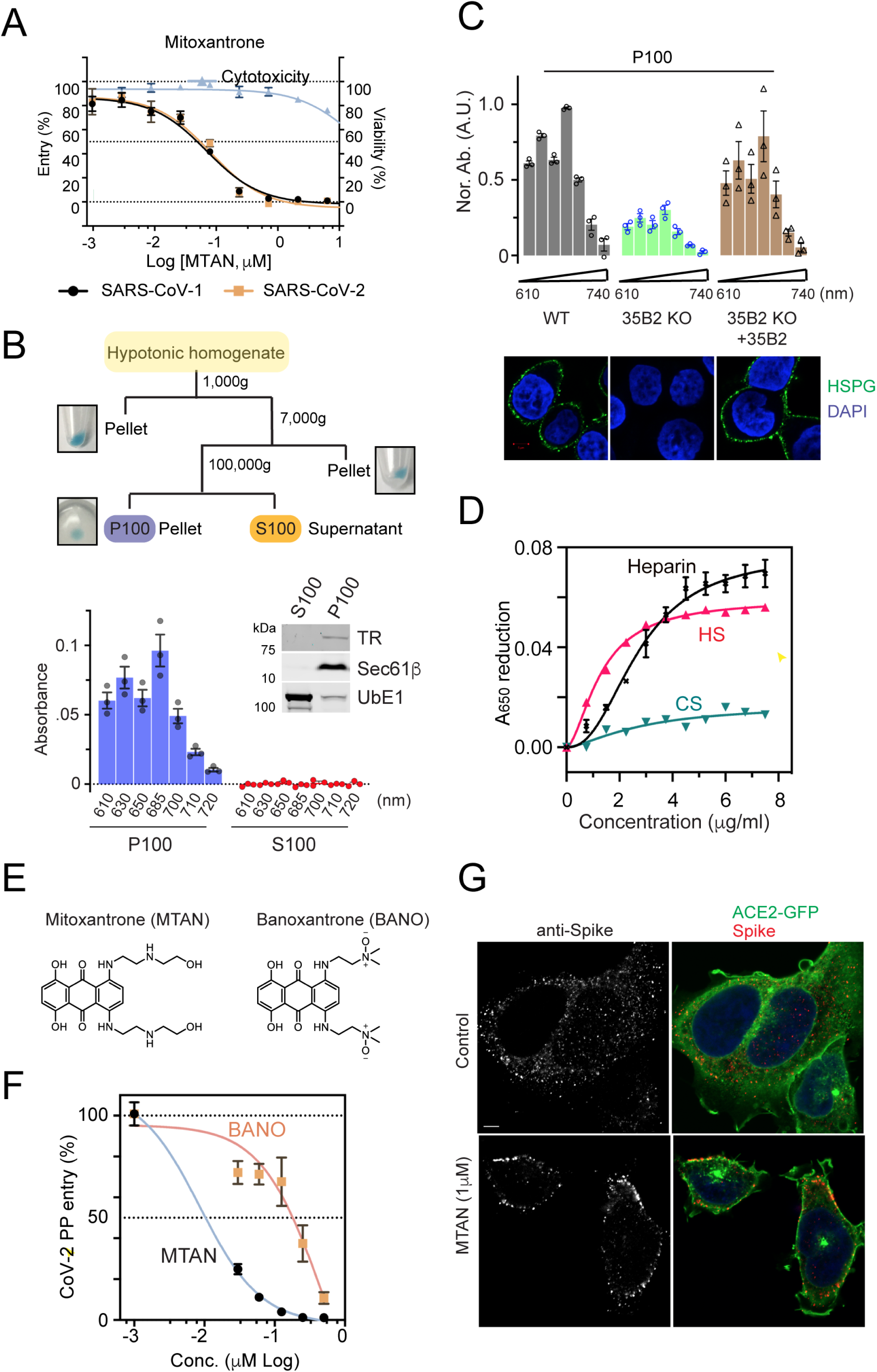
Mitoxantrone targets HS directly to inhibit SARS coronavirus entry. **(A)** Mitoxantrone inhibits Spike-mediated entry of pseudoviral particles (PP). ACE2-GFP HEK293 cells were transduced with SARS-CoV and SARS-CoV-2 PP for 48 h in the presence of Mitoxantrone as indicated. Error bars indicate SEM, N=4. Cells treated without the virus were used to control drug cytotoxicity. **(B)** The subcellular distribution of Mitoxantrone. The scheme illustrates the experimental procedure. HEK293T cells were treated with 5 μM Mitoxantrone for 30 min at 37 °C before fractionation. The membrane pellet (P100) and the cytosol supernatant (S100) fractions were analyzed by immunoblotting and by spectrometry at the indicated wavelengths. Error bars indicate SEM. N=3. **(C)** The membrane association of Mitoxantrone requires *SLC35B2 (35B2)*. Cells of the indicated genotypes were treated with 5 μM Mitoxantrone for 30 min at 37 °C and then fractionated as in B. The absorbance of the P100 fractions was measured and the control sample at 680 nm was normalized to 1. Bottom panels show the cells stained with DAPI (blue) and GFP+ (green) to label DNA and HS (Green), respectively. Error bars indicate SEM. N=3. **(D)** Mitoxantrone has a higher affinity for heparin and HS than chondroitin sulfate (CS). A_650_ of Mitoxantrone (5 μM) mixed with the indicated oligosaccharides was determined. The decrease in absorbance after the addition of the oligosaccharides was plotted. Error bars indicate SEM. N=2. **(E)** The chemical structures and absorbance spectra of Mitoxantrone (MTAN) and Banoxantrone (BANO). **(F)** Banoxantrone (BANO) has reduced antiviral activity. ACE2-GFP HEK293T cells were incubated with SARS-CoV-2 in the presence of the indicated drugs for 24 h before measuring the luciferase activity. Error bars indicate SEM. N=4. **(G)** Mitoxantrone inhibits SARS-CoV-2 cell entry. ACE2-GFP HEK293T cells were pretreated with DMSO as a control or Mitoxantrone for 30 min before incubation with SARS-CoV-2 PP. Cells were fixed 3 h later and stained with anti-Spike antibodies (red) and DAPI (blue).

Because Mitoxantrone has absorbance peaks at 620 nm and 685 nm (Fig. S4A). we used this spectral property to trace its localization in drug-treated cells, which hinted at its cellular target. To this end, we performed subcellular fractionation using Mitoxantrone-treated cells to obtain the nucleus (1,000g pellet), mitochondria-enriched heavy membrane (7,000g pellet), light membrane (100,000g pellet), and cytosol (100,000g supernatant) fractions (Figure 5B top panel). Although blue color was seen in every pellet fraction, Mitoxantrone in the nucleus and heavy membrane fractions was resistant to extraction by a buffer containing the non-ionic detergent NP40 or 1% SDS, probably due to tight association with DNA. By contrast, Mitoxantrone in the light membranes (containing the plasma membrane and endoplasmic reticulum as demonstrated by immunoblotting, middle panel) could be readily released by an NP40-containing buffer and detected by a spectrometer (Figure 5B). No Mitoxantrone was detected in the cytosol fraction. Thus, in addition to DNA, Mitoxantrone also binds to cell membranes.

Several lines of evidence suggest that Mitoxantrone targets the cell surface HS directly. First, the amount of P100-associated Mitoxantrone from *SLC35B2* deficient cells (lacking HS) was significantly lower than that of wild-type cells, which could be rescued by re-expression of SLC35B2 (Figure 5C). Additionally, as mentioned above, Mitoxantrone readily bound to heparin, a glycan structurally related to HS (Figure 3F, Fig. S4B). Because the interaction of heparin with Mitoxantrone lowered its absorption spectrum in solution (Fig. S4C), we used this property to further determine the interaction of Mitoxantrone with other related glycans. As expected, HS bound to Mitoxantrone similarly as heparin, but chondroitin sulfate (CS) did not bind to Mitoxantrone significantly (Figure 5D). Moreover, Banoxantrone, an anti-cancer drug structurally related to Mitoxantrone did not show significant interaction with either heparin or HS (Figure 5E, Fig. S4D, E). These results suggest a specific and direct interaction between Mitoxantrone and HS, which explains the SLC35B2-dependent association with the cell membranes. Importantly, compared to Mitoxantrone, Banoxantrone was a much weaker SARS-CoV-2 PP entry inhibitor (Figure 5F), suggesting a crucial role for heparin/HS binding in the anti-viral activity of Mitoxantrone.

Although Mitoxantrone binds HS directly, it did not disrupt the binding of SARS-CoV-2 PP to ACE2-GFP-expressing cells. Instead, when cells pretreated with Mitoxantrone were exposed to SARS-CoV-2 PP, most viral particles remained bound to the plasma membrane as revealed by immunostaining with an anti-Spike antibody (Figure 5G). By contrast, in control cells, Spike antibodies detected mostly intracellular speckles, probably representing endocytosed viral particles (Figure 5G). This result suggests that Mitoxantrone and Spike bind HS in different modes, but the binding of Mitoxantrone to HS alters its function to block viral entry.

Surprisingly, while weaker inhibitors like BNTX rescued the CPE of SARS-CoV-2 in Vero E6 cells, Mitoxantrone did not show any activity in this assay. This might be due to cytotoxicity linked to the DNA binding off-target property of Mitoxantrone. Consistent with this notion, Vero E6 cells were more sensitive to Mitoxantrone-induced cell death (Fig. S4F vs. Figure 5A). In this regard, it is compelling to see that the dose-dependent toxicity profiles of Mitoxantrone and Banoxantrone were indistinguishable (Fig. S4F), which suggests the possibility of generating Mitoxantrone derivatives with reduced cytotoxicity but similar antiviral activity.

### A combination regimen optimally targeting the HS-assisted viral entry

We next used gene expression profiling to further define the action of Tilorone, Raloxifene, and Piceatannol in comparison to Mitoxantrone. We treated HEK293T cells with these drugs at a dose that inhibited Spike-mediated viral entry. Differential gene expression analyses showed that Mitoxantrone had the most significant impact on gene expression, probably due to its strong DNA binding activity (Figure 6A). By contrast, Tilorone had the smallest effect, altering a limited number of genes by only small changes (Fig. S5A-D). Cluster analysis showed that the gene expression signature associated with Raloxifene treatment was more similar to that of Piceatannol-treated cells than to Tilorone-treated cells (Figure 6A), consistent with the observation that Raloxifene and Piceatannol but not Tilorone interact with heparin.

**Figure 6.**
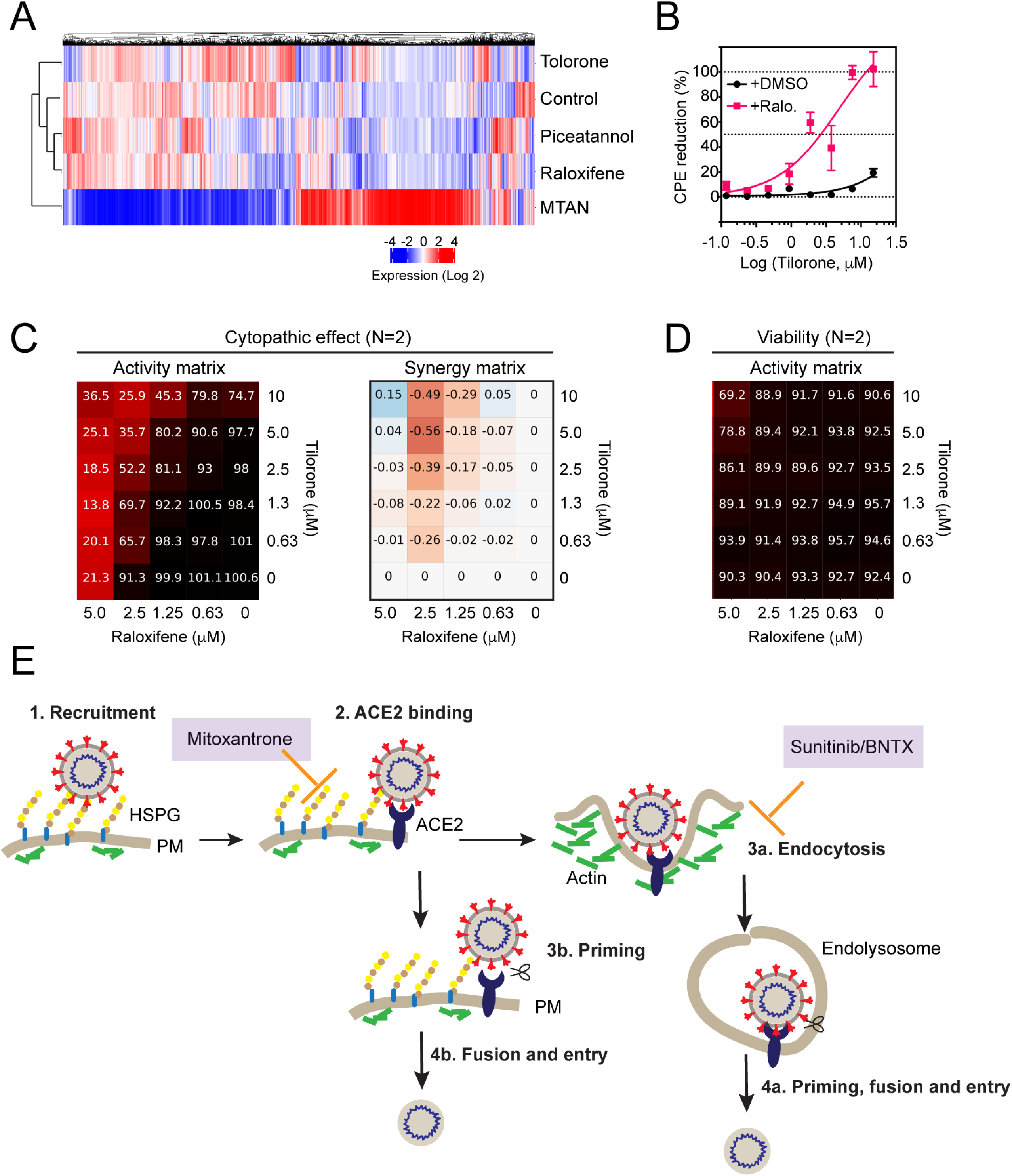
A combination regimen that optimally targets HS-dependent viral entry. (**A**) A heatmap showing the gene expression profiles of untreated HEK293T cells or cells treated with the indicated drugs at 10 μM for 6 h (Mitoxantrone 5 μM). (**B-D**) Drug combination studies show a synergistic CPE rescue effect by Tilorone and Raloxifene (Ralo.). (B) Vero E6 cells treated with Tilorone at different concentrations with (1.5 μM) or without Raloxifene were infected with SARS-CoV-2 for 72 h before viability test. Error bars indicate SEM. N=2. (C, D) Matrix blocks for Tilorone plus Raloxifene in the CPE and cell viability assays. Activity blocks show normalized levels of viral toxicity (C) or normalized cell viability (no virus) (D). Red labels in the synergy matrix indicate positive synergism whereas blue labels suggest antagonism. (**E**) A SARS-CoV-2 entry map with the drug targeting sites identified in this study. PM, plasma membrane.

Conceptually, it might be possible to combine drugs targeting distinct steps in the HS-assisted entry pathway to produce synergized anti-viral activity. We tested the combination of Raloxifene with Tilorone because the latter has been used as an anti-viral agent and because either Tilorone or Raloxifene by itself modestly rescued SARS-CoV-2-induced CPE in Vero E6 cells (Fig. S5E, F). Strikingly, co-treatment with 1.5 μM Raloxifene reduced IC_50_ of Tilorone by ∼10-fold in the CPE assay (Figure 6B). The synergistic inhibition of viral CPE was further confirmed by a matrix test, in which Tilorone was tested systematically with different concentrations of Raloxifene (Figure 6C). Importantly, no significant cytotoxicity was observed in combined treatment even at highest concentrations (Figure 6D).

## Discussion

It is known that ACE2-mediated SARS-CoV and CoV-2 entry is controlled by a protease-dependent priming step ^14-16^. Evidence presented in this study supports an additional layer of regulation imposed by the cell surface HS. The cell surface HS proteoglycans comprise of two protein families (6 Glypicans and 4 Syndecans) ^32^. Proteins of these two families share no sequence similarity, but they all bear negatively charged HS polymers that can promote cell interactions with a variety of endocytic ligands ^20, 21^. Our data together with recent preprints in BioRxiv support a model in which HS glycans help to recruit SARS-CoV-2 to the cell surface, which increases its local concentration for effectively engaging ACE2 ^43, 44^ (Figure 6E). Interestingly, some other coronaviruses also use a similar two-receptor mechanism for cell entry. For example, some α-Coronaviruses first attach themselves to sialoglycans on the cell surface before binding to proteinaceous receptors ^45, 46^.

How do HS glycans interact with Spike? Two recent studies posted at BioRxiv modeled the interactions of heparin with Spike using either full length Spike or just the RBD domain ^29, 44^. While several potential binding sites were suggested on full length Spike, a more sensible model was generated when the RBD domain was used. This model suggests a long positive charge-enriched binding groove that accommodates a chain of heparin/HS. This model, if confirmed, is consistent with the observed salt sensitive heparin-Spike interaction.

How Mitoxantrone inhibits SARS-CoV and CoV-2 entry remains to be elucidated. It is known that Spike in the prefusion state adopts distinct conformations with the RBD domain either in the up or down position ^44, 47^, and the interaction of Spike with ACE2 requires the RBD-up conformation ^11, 48-50^. By contrast, the proposed HS binding model is compatible with either the RBD-up or down conformation ^44^. Thus, it is possible that HS binding occurs on Spike with flexible RBD positions, and this conformational plasticity may facilitate downstream steps such as ACE2-dependent endocytosis and/or protease-mediated priming ^51^. Mitoxantrone might influence the way HS interacts with the Spike and thus inhibits viral entry.

In addition to Mitoxantrone, we uncover several other drugs capable of targeting HS and HS-dependent endocytosis. The fact that many of these drugs also mitigate Spike-dependent viral entry strongly corroborates the idea that ACE2-mediated endocytosis serves a major role in SARS-CoV-2 cell entry. The identified drugs have a relatively limited impact on endocytosis as they affect neither clathrin-mediated transferrin uptake nor clathrin-independent uptake of CTB. Thus, these drugs are expected to be less toxic than previously reported pan-endocytosis inhibitors ^13^. Among the drug identified, Sunitinib and BNTX disrupt actin dynamics (Figure 7E), suggesting the actin network as a previously unknown regulator for SARS-CoV and CoV-2 entry. These viruses may enter cells using receptor-mediated macropinocytosis, which depends on actin and has been known for its role in in viral pathogen uptake ^52^.

Although our study focuses on SARS-CoV and CoV-2, drugs targeting HS and HS-dependent endocytosis are expected to have a broad antiviral activity. Consistent with this view, Tilorone has long been recognized as a general antiviral agent. An early study in mice suggested that Tilorone administrated orally might upregulate interferon production ^53^, but our gene expression study did not reveal significant changes in the expression of interferon by Tirolone treatment. Instead, the demonstration of Tilorone as an inhibitor of HS-dependent endocytosis offers an alternative explanation for the reported antiviral function. Importantly, our study also suggests that Raloxifene, a heparin/HS-binding drug can enhance the anti-viral activity of Tilorone. Raloxifene is a selective estrogen receptor modulator used to treat osteoporosis and to prevent breast cancer in postmenopausal women (drugbank.ca/drugs/DB00481) ^54^. Both Tilorone and Raloxifene are well tolerated and can be conveniently administrated orally. Thus, whether this combination can be clinically effective against SARS-CoV-2 infection deserves additional testing in animal and other anti-viral models. In summary, our study not only establishes HS-dependent viral entry as a novel drug target for COVID-19, but also reveals candidate drugs that disrupt this critical event in the SARS-CoV-2 life cycle.

## Methods

### Chemicals and reagents

The initial screening library LOPAC^R1280^ was purchased from Sigma. The chemicals for the follow-up studies were purchased as indicated in the table below. Standard quality control by HPLC was conducted before the drugs were used in the study. pcDNA3.1-SARS-CoV2-Spike plasmid was obtained from BEI resource ^11^. Other chemicals and reagents were listed in the table below.

### Cell line, transfection, lentivirus production, and infection

HEK293T cells stably expressing human ACE2 tagged with GFP (ACE2-GFP) were generated by transfecting cells with pCMV-ACE2-GFP (Covex), and stable clones were hand-picked after neomycin (1 mg/ml) selection for 1 week. Two ACE2-GFP cell lines using either HEK293 or HEK293T as the parental line were independently generated and used at NCATS and NIDDK, respectively. U2OS cells stably expressing Clathrin light chain-mCherry and Tractin-EGFP and the CRISPRv2 construct expressing sgRNA targeting *SLC35B2* were described previously ^19^. Calu-3 cells were purchased from ATCC and maintained in MEM with 10% fetal bovine serum. sgRNA-expressing lentiviruses were produced by transfecting 1 million 293FT cells (Thermo Fisher Scientific) in a 3.5 cm dish with 0.4 μg pVSV-G, 0.6 μg psPAX2, and 0.8 μg CRISPRv2-sgRNA. Transfected cells were incubated with 3 ml fresh DMEM medium for 72 h before viruses were harvested.

### α-Synuclein fibrils uptake assay and drug screen

Fluorescence dye labeled α-Syn preformed fibrils were generated as previously described ^19^. siRNA-mediated gene silencing was performed by lipofectamine RNAiMAX (Invitrogen) mediated transfection following the instruction from the manufacture. HEK293T cells were seeded in white 1,536-well microplates that have transparent bottom (Greiner BioOne) at 2,000 cells/well in 2 μL media and incubated at 37 °C overnight (∼16 h). pHrodo red-labeled α-Syn fibrils were added at 400 nM final concentration to each well by a dispenser. After 1 h incubation, compounds from the LOPAC^R1280^ library (Sigma) were titrated 1:3 in DMSO and dispensed via pintool at 23 nl/well to the assay plates. After 24 hours of incubation, the fluorescence intensity of pHrodo red was measured by a CLARIOstar Plus plate reader (BMG Labtech). Data were normalized using cells containing 400 nM pHrodo red-labeled Syn fibrils as 100% and medium containing 400 nM preformed fibrils-pHrodo red as 0%.

### SARS-CoV and SARS-CoV-2 pseudotyped particles (PP)

SARS-CoV-S, SARS-CoV2-S, and ΔEnv (lack the Spike protein) pseudotyped particles (PP) were custom manufactured by the Codex Biosolutions (Gaithersburg, MD) following previously reported methods using a murine leukemia virus (MLV) pseudotyping system ^55, 56^. The SARS-CoV2-S construct with Wuhan-Hu-1 sequence (BEI #NR-52420) was C-terminally truncated by 19 amino acids to reduce ER retention for pseudotyping ^12^.

### PP entry assay in the 1536-well format

HEK293-ACE2 cells seeded in white, solid bottom 1536-well microplates (Greiner BioOne) at 2,000 cells/well in 2 μL medium were incubated at 37 °C with 5% CO2 overnight (∼16 h). Compounds were titrated 1:3 in DMSO and dispensed via pintool at 23 nl/well to the assay plates. Cells were incubated with compounds for 1 h at 37 °C with 5% CO_2_ before 2 μl/well of PP were added. The plates were then spinoculated by centrifugation at 1,500 rpm (453 x g) for 45 min and incubated for 48 h at 37 °C 5% CO2 to allow cell entry of PP and the expression of luciferase. After the incubation, the supernatant was removed with gentle centrifugation using a Blue Washer (BlueCat Bio). Then 4 μL/well of Bright-Glo luciferase detection reagent (Promega) was added to assay plates and incubated for 5 min at room temperature. The luminescence signal was measured using a PHERAStar plate reader (BMG Labtech). Data were normalized with wells containing PP as 100% and wells containing control ΔEnv PP as 0%. Experiments shown in Figure 4A-C, Fig. S3A, Figure 5A were conducted in this format at NCATS.

### PP entry assay in the 96-well format

HEK293T-ACE2-GFP cells were seeded in white, transparent bottom 96-well microplates (Thermo Fisher Scientific) at 20,000 cells per well in 100 μl growth medium and incubated at 37 °C with 5% CO2 overnight (∼16 h). The growth medium was carefully removed and 50 μl PP or PP containing compounds were added into each well. The plates were then spinoculated by centrifugation at 1500 rpm (453 x g) for 45 min and incubated for 24h (48 h for Calu-3 cells) at 37 °C 5% CO_2_ to allow cell entry of PP and the expression of luciferase. After incubation, the supernatant was carefully removed. Then 50 μl/well of Bright-Glo luciferase detection reagent (Promega) was added to assay plates and incubated for 5 min at room temperature. The luminescence signal was measured by a Victor 1420 plate reader (PerkinElmer). For ACE2-GFP cells, the GFP signal was also determined by the plate reader. Data were normalized with wells containing PP but no compound as 100%, and wells mock-treated with phosphate buffer saline (PBS) as 0%, and the ratio of luciferase to the corresponding GFP intensity was calculated. Experiments shown in Figure 1B, D, Figure 2B, C, Figure 5F were done following this protocol at NIDDK.

### ATP content cytotoxicity assay in the 1536-well format

HEK293-ACE2 cells were seeded in white, solid bottom 1,536-well microplates (Greiner BioOne) at 2,000 cells/well in 2 μl medium and incubated at 37 °C with 5% CO2 overnight (∼16 h). Compounds were titrated 1:3 in DMSO and dispensed via pintool at 23 nl/well to assay plates. Cells were incubated for 1 h at 37 °C 5% CO2 before 2 μl/well of media was added. The plates were then incubated at 37 °C for 48 h at 37C 5% CO2. After incubation, 4 μl/well of ATPLite (PerkinElmer) was added to assay plates and incubated for 15 min at room temperature. The luminescence signal was measured using a Viewlux plate reader (PerkinElmer). Data were normalized with wells containing cells as 100%, and wells containing media only as 0%.

### ATP content cytotoxicity assay in the 96-well format

HEK293T-ACE2-GFP cells were seeded in white, transparent bottom 96-well microplate (Thermo Fisher Scientific) at 20,000 cells per well in 100 μl/well growth medium and incubated at 37 °C with 5% CO2 overnight (∼16 h). The growth medium was carefully removed and 100 μl medium with compounds was added into each well. The plates were then incubated at 37 °C for 24 h (48 h for Calu3 cells) at 37 °C 5% CO2. After incubation, 50 μL/well of ATPLite (PerkinElmer) was added to assay plates and incubated for 15 min at room temperature. The luminescence signal was measured using a Victor plate reader (PerkinElmer). Data were normalized with wells containing cells but no compound as 100%, and wells containing media only as 0 %.

### SARS-CoV-2 cytopathic effect (CPE) assay

SARS-CoV-2 CPE assay was conducted at the Southern Research Institute (Birmingham, AL) as fee-for-service. Briefly, compounds were titrated in DMSO and acoustically dispensed into 384-well assay plates at 60 nl/well. Cell culture medium (MEM, 1% Pen/Strep/GlutaMax, 1% HEPES, 2% HI FBS) was dispensed at 5 μl/well into assay plates and incubated at room temperature. Vero E6 (selected for high ACE2 expression) was inoculated with SARS CoV-2 (USA_WA1/2020) at 0.002 M.O.I. in media and quickly dispensed into assay plates at 4,000 cells/well in 25 μl volume. Assay plates were incubated for 72 h at 37 °C, 5% CO_2_, 90% humidity. Then, 30 μl/well of CellTiter-Glo (Promega) was dispensed, incubated for 10 min at room temperature, and the luminescence signal was read on an EnVision plate reader (PerkinElmer). An ATP content cytotoxicity assay was conducted with the same protocol as CPE assay except that SARS-CoV-2 virus was omitted from the incubation.

### GFP+ purification and HSPG staining

His-tagged GFP+ was reported previously ^19^. To stain the cell surface HSPG by GFP+, we incubated cells with 200 nM GPF+ in the growth medium on ice for 15min. Cells were then fixed and stained with DAPI to reveal the nucleus.

### Endocytosis and viral binding assays

To test the effect of the compounds on DNA uptake, we seeded HEK293T cells previously infected with a YFP-expressing retrovirus at 0.2 x 10^6^/well in a poly-lysine D coated 12 well plate. 24 h later, cells were treated with the chemicals for 30 min before a transfection mixture containing 0.2 μg mCherry-expressing plasmid and 0.6 μl TransIT293 (Mirus) was added to the cells. Cells were incubated with the DNA for 4 h, after which the medium was replenished. The cells were further grown for 48 h. Cells were then lysed in NP40 lysis buffer and the fluorescence intensity in cleared lysates was measured by a Fluoromax3 fluorometer (Horiba).

To test the effect of compounds on VSVG-coated lentivirus uptake/infection, HEK293T cells expressing YFP were seeded at 0.2×10^6^/well in a poly-lysine D coated 12 well plate. After 24 h, cells were treated with compounds for 30 min and then infected with a mCherry-expressing lentivirus at M.O.I. of ∼ 2.0 in the presence of the compounds for 6 h. The virus and compounds were removed, and cells were further grown in fresh medium for 48 h before fluorescence measurement.

To test the effect of compounds on GFP+ or transferrin uptake, HEK293T cells were seeded at 25,000 cells per well in an 8 well chamber (ibidi). 24 h later, cells were treated with compounds for 30 min before the addition of GFP+ (200 nM) or transferrin (Thermo Fisher Scientific) (50 μg/ml). Cells were further incubated at 37 °C for 4 h before fixation and confocal imaging.

To detect the binding of SARS-CoV-2 to the cell surface, HEK293T-ACE2-GFP cells seeded in 24 well plates that had been coated with fibronectin were treated with 50 μl virus per well at 4 °C for 1 h with centrifugation at 1500 rpm (453 x g) for 60 min. Cells were carefully washed with ice-cold PBS to remove unbound virus and then lysed in a NP40 lysis buffer containing 0.5 % Igepal, 20 mM Tris pH 7.4, 150 mM Sodium Chloride, 2 mM Magnesium Chloride, 0.5 mM EDTA, 1 mM DTT, and a protease inhibitor cocktail. The cell extracts cleared by centrifugation (16,000 x g 5 min) were analyzed by immunoblotting.

### UPLC-MS/MS assay

Ultra-performance liquid chromatography-tandem mass spectrometry (UPLC-MS/MS) methods were developed and optimized to determine compounds’ concentrations in the in vitro samples. Mass spectrometric analysis was performed on a Waters Xevo TQ-S triple quadrupole instrument using electrospray ionization in positive mode (Mitoxantrone, BNTX, Tilorone and Sunitinib) and negative mode (Raloxifene, Piceatannol, Exifone, K114) with the selected reaction monitoring. The separation of test compounds was performed on an Acquity BEH C_18_ column (50 ⨯ 2.1 mm, 1.7 μ) using a Waters Acquity UPLC system with 0.6 mL/min flow rate and 2 minute gradient elution. The mobile phases were 0.1% formic acid in water and 0.1% formic acid in acetonitrile. The calibration standards (0.100 – 100 μM) and quality control samples were prepared in 50% acetonitrile/water with 0.1% formic acid and PBS buffer. Aliquots of 10 μL samples were mixed with 200 μL internal standard in acetonitrile to precipitate proteins in a 96-well plate. 0.5 μL supernatant was injected for the UPLC-MS/MS analysis. Data were analyzed using MassLynx V4.1 (Waters Corp., Milford, MA).

### qRT-PCR analysis of knockout or knockdown cells

Total RNA was extracted from 3 million HEK293T?cells using TriPure reagent (Roche) and purified using RNeasy MinElute Cleanup Kit (Promega) following the standard protocols. The RNA concentration was measured by Nanodrop 2000 UV spectrophotometer, and 1μg total RNA was converted to cDNA using the iScript Reverse Transcription Supermix (BioRad) system. 1μL cDNA was used to perform qPCR using SsoAdvanced SYBR Green supermix kit (BioRad) on a CFX96 machine (BioRad). Data were analyzed using BioRad CFX Manager 3.0 software. GADPH was used as a reference gene for the quantification of gene expression levels. Primers used for qRT-PCR were listed in the previous study ^19^. For the RNAseq study, cells were treated independently for 6 h with each drug three times and 6 untreated control samples were included. The treatment conditions are Mitoxantrone 5 μM, Tilorone 10 μM, Raloxifene 10 μM, Piceatannol 10 μM. RNA isolated from control or drug-treated cells (10 μg/sample) were processed by Novagene USA.

### Membrane fractionation and heparin Sepharose pulldown

To fractionate cells, ∼15 million cells were treated with 5 μM Mitoxantrone or Banoxantrone for 30 min at 37 °C. Cells were harvested and washed with ice-cold PBS. Cells were then resuspended and incubated in 900 μl of a hypotonic buffer (50 mM HEPES, pH 7.3, 25 mM potassium acetate) containing a protease inhibitor cocktail. Cells were homogenized in a Dounce homogenizer with a tight pestle and then subject to differential centrifugation at 1,000 g for 5 min, 7,000 x g for 10min, and 100,000 x g for 20 min. The P100 membrane pellet was resuspended in 20 μl NP40 lysis buffer containing 20 mM Tris pH 7.4, 0.5% NP40, 150 mM Sodium Chloride, 2 mM Magnesium Chloride. The absorbance in the cleared membrane extract and the S100 fraction was measured by a NanoDrop 2000 spectrometer. Note that no absorbance was detected for the S100 fraction even when the samples were measured by a conventional spectrometer with a 10 x light path.

To determine the binding of Mitoxantrone to heparin-coated Sepharose, the compound was diluted to 50 μM in PBS and then incubated with PBS-washed heparin beads or as a control with Sepharose. After brief mixing, the beads were sedimented by centrifugation. The supernatant fractions were analyzed by NanoDrop 2000 for light absorption at the wavelengths specified in the figure legends.

The pulldown of Spike by heparin beads were done by incubating 600ng Spike with pre-washed heparin Sepharose in 25mM Tris pH 7.3, 2mM KCl, 1mM MgCl_2_, 0.05% Igepal with or without 150mM NaCl. To test the effect of Mitoxantrone on heparin-Spike interaction, 30 μl heparin beads were washed with PBS, treated with 200 μl Mitoxantrone (100 μM) for 5 min at room temperature or as a control remained untreated. Unbound Mitoxantrone was removed after centrifugation. The beads were then incubated with 600 ng purified Spike protein in 300 μl PBS containing 0.05% NP40 and 1mM DTT. The reaction was incubated at room temperature for 30 min. The beads were washed two times with PBS and eluted with 1 x Laemmli buffer for SDS-PAGE and immunoblotting analysis.

### Drug synergy analysis

CPE and Toxicity were normalized using independent control wells (DMSO ± virus) on each plate, so activity values were not strictly bounded between [0, 100]. For CPE assay, DMSO+virus was treated as the neutral control, whereas DMSO-only (no virus) served as the positive control. Then normalized CPE = 1 - (x - neutralCtrl) / (positiveCtrl - neutralCtrl) × 100%. For viability assay, DMSO-only was used as the neutral control and media-only wells (no cell) as the negative control. Normalized viability = (x - negativeCtrl) / (neutralCtrl - negativeCtrl) × 100%. Synergism and antagonism from a 6 × 6 block were evaluated using the <u>h</u>ighest <u>s</u>ingle <u>a</u>gent model (HSA). Given a dose combination A_conc1_ + B_conc2_,

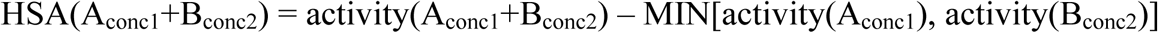

Therefore, we have

Synergism: HSA(*) < 0 (value from combination is lower than the best single agent)

Antagonism: HSA(*) > 0 (value from combination is greater than the best single agent)

Additivity: HSA(*) = 0 (value from combination equals to best single agent)

The overall HSA synergism (HSA_sum_) given a 6 × 6 block was calculated as the sum of HSA(A_conc*_+B_conc*_) across the non-toxic dose combinations (defined as viability activity > 80). We used empirical cutoff HSA_sum_ < −100 to call a synergistic combination. The highest concentration of Raloxifene was excluded from the figure because it alone induced significant cytotoxicity.

### Image processing and statistical analyses

Confocal images were processed using the Zeiss Zen software. To measure fluorescence intensity, we used the Fiji software. Images were converted to individual channels and region of interest was drawn for measurement. Statistical analyses were performed using either Excel or GraphPad Prism 8. Data are presented as mean ± SEM, which was calculated by GraphPad Prism 8. *p*-values were calculated by Student’s t-test using Excel. None linear curve fitting and IC_50_ calculation was done with GraphPad Prism 8 using the inhibitor response 3 variable model or the exponential decay model. Images were prepared with Adobe Photoshop and assembled in Adobe Illustrator. All experiments presented were repeated at least twice independently except for the data from the Southern Research Institute, which was performed as fee-for-service in 2 duplicates. Data processing and reporting are adherent to the community standards.

## Data Availability

All data, associated protocols, methods, and sources of materials can be accessed in the main text or supplementary information. The analysis code for drug synergy study is available at NCATS github. The mRNA sequencing data has been deposited to NCBI Sequence Read Archive. The accession ID is: PRJNA645209.

### Acknowledgements

We thank Susan Buchanan, Martin Gellert (NIDDK), and Liqiang Chen (University of Minnesota) for critical reading of the manuscript. The plasmids and protocols for pseudotyped particle generation were kind gifts from Dr. Gary Whittaker at Cornell University. The work is supported by the Intramural Research Program of the National Institute of Diabetes, Digestive & Kidney Diseases and of the National Center for Advancing Translational Sciences in the National Institutes of Health.

## Author contribution

Q. Zhang, M. Swaroop, W. Zheng, and Ye, Y. designed the drug screen, Q. Zhang, C. Chen, Y. Xu, W. Zheng, and Ye Y. conceived and designed the study, A.Q. Wang, X. Xu, and C. Chen designed the mass spectrometry experiment, Q. Zhang, C. Chen, Y. Xu, W. Zheng, L. Chen, A.Q. Wang, and Ye Y. analyzed the data. Q. Zhang, M. Swaroop, M. Pradhan, M. Xu, L. Wang, J. Lee, M. Shen, Z. Luo, W. Huang, A.Q. Wang, X. Xu, N. Hagen, and Y. Ye conducted the experiments. Y. Ye wrote the manuscript. All authors contribute to the editing of the manuscript.

## List of supplementary materials

**Supplementary Table S1.**

| Reagents and Materials | Source | Catalogue No. |
| --- | --- | --- |
| pHrodo™ Red, succinimidyl ester (pHrodo™ Red, SE) | ThermoFisher Scientific | Cat# P36600 |
| Banoxantrone dihydrochloride | Sigma | Cat# SML1854 |
| Heparin sodium salt from porcine intestinal mucosa | Sigma | Cat# H3393 |
| Heparan sulfate sodium salt | Biosynth Carbosynth | YH30121 |
| Chondroitin sulfate sodium salt | Biosynth Carbosynth | YH04273 |
| SARS-CoV-S and SARS-CoV2-S Pseudotyped particles | Codex Biosolutions (Gaithersburg, MD) | Contracted custom production |
| ATPLite | PerkinElmer | Cat# 6016736 |
| Bright-Glo Luciferase kit | Promega | Cat# E2620 |
| CellTiter-Glo cell viability kit | Promega | Cat# G7572 |
| iScript™ Reverse Transcription Supermix for RT-qPCR | BioRad | Cat# 1708840 |
| SsoAdvanced Universal SYBR Green Supermix | BioRad | Cat# 1725271 |
| Imaging chamber | iBidi | Cat# 80426 |
| TriPure reagent | Sigma | Cat# 11667157001 |
| RNeasy MinElute Cleanup Kit | Qiagen | Cat# 74024 |
| Latrunculin A | TOCRIS | Cat# 3973 |
| NCGC00015693-09 | Microsource | 01503278 |
| NCGC00015889-11 | Microsource | 01505622 |
| NCGC00024857-02 | BIOMOL | AC-304 |
| NCGC00094226-07 | SigmaAldrich | Lopac-P-0453 |
| NCGC00094226-13 | Selleck | S3026 |
| NCGC00095196-06 | Microsource | 02300009 |
| NCGC00164631-05 | Sequoia | SRP01785s |
| NCGC00186034-01 | SigmaAldrich | Lopac-K-1015 |
| NCGC00186034-03 | SIGMA | K1015 |
| NCGC00249050-02 | GVK | FFS-12-73-NCI-3078 |
| Spike S1_S2 | Sino Biological | 40589-V27B-B |
| pLV-mCherry | A gift from Pantelis Tsoulfas (Addgene #36084) |  |
| pcDNA3.1-SARS-CoV2-Spike | BEI resources | NR-52420 |
| SARS-CoV-2 S antibody | GeneTex | Cat# GTX632604 |

Figures S1-S6

Movies S1-S6

## Supplementary movies

**Movie 1**

A U2OS cell expressing Tractin-EGFP.

**Movie 2**

The cell shown in movie 1 was imaged after treatment with Sunitinib 5 μM for 15 min. Note the increased assembly of actin filaments causes new filopodia formation on the cell surface.

**Movie 3**

A Tractin-EGFP-expressing U2OS cell was imaged after treatment with Sunitinib 5 μM for 60 min.

**Movie 4**

The same Tractin-EGFP-expressing U2OS cell in movie 3 was imaged at a different confocal plan where the stress fibers are located.

**Movie 5**

A U2OS cell expressing Tractin-EGFP.

**Movie 6**

The cell shown in movie 5 was imaged after treatment with BNTX 10 μM for 5min.

## References

1 Zhou P, Yang XL, Wang XG et al. A pneumonia outbreak associated with a new coronavirus of probable bat origin. Nature 2020; 579:270–273.

2 Zhu N, Zhang D, Wang W et al. A Novel Coronavirus from Patients with Pneumonia in China, 2019. N Engl J Med 2020; 382:727–733.

3 Holmes KV. SARS-associated coronavirus. N Engl J Med 2003; 348:1948–1951.

4 Marsh M, Helenius A. Virus entry: open sesame. Cell 2006; 124:729–740.

5 Tortorici MA, Veesler D. Structural insights into coronavirus entry. Adv Virus Res 2019; 105:93–116.

6 Wang H, Yang P, Liu K et al. SARS coronavirus entry into host cells through a novel clathrin- and caveolae-independent endocytic pathway. Cell Res 2008; 18:290–301.

7 Burkard C, Verheije MH, Wicht O et al. Coronavirus cell entry occurs through the endo-/lysosomal pathway in a proteolysis-dependent manner. PLoS Pathog 2014; 10:e1004502.

8 Belouzard S, Millet JK, Licitra BN, Whittaker GR. Mechanisms of coronavirus cell entry mediated by the viral spike protein. Viruses 2012; 4:1011–1033.

9 Inoue Y, Tanaka N, Tanaka Y et al. Clathrin-dependent entry of severe acute respiratory syndrome coronavirus into target cells expressing ACE2 with the cytoplasmic tail deleted. J Virol 2007; 81:8722–8729.

10 Glebov OO. Understanding SARS-CoV-2 endocytosis for COVID-19 drug repurposing. FEBS J 2020.

11 Walls AC, Park YJ, Tortorici MA, Wall A, McGuire AT, Veesler D. Structure, Function, and Antigenicity of the SARS-CoV-2 Spike Glycoprotein. Cell 2020; 181:281–292 e286.

12 Ou X, Liu Y, Lei X et al. Characterization of spike glycoprotein of SARS-CoV-2 on virus entry and its immune cross-reactivity with SARS-CoV. Nat Commun 2020; 11:1620.

13 Kang YL, Chou YY, Rothlauf PW et al. Inhibition of PIKfyve kinase prevents infection by EBOV and SARS-CoV-2. bioRxiv 2020.

14 Hoffmann M, Kleine-Weber H, Schroeder S et al. SARS-CoV-2 Cell Entry Depends on ACE2 and TMPRSS2 and Is Blocked by a Clinically Proven Protease Inhibitor. Cell 2020; 181:271–280 e278.

15 Hoffmann M, Kleine-Weber H, Pohlmann S. A Multibasic Cleavage Site in the Spike Protein of SARS-CoV-2 Is Essential for Infection of Human Lung Cells. Mol Cell 2020; 78:779–784 e775.

16 Shang J, Wan Y, Luo C et al. Cell entry mechanisms of SARS-CoV-2. Proc Natl Acad Sci U S A 2020; 117:11727–11734.

17 Millet JK, Whittaker GR. Physiological and molecular triggers for SARS-CoV membrane fusion and entry into host cells. Virology 2018; 517:3–8.

18 Zang R, Gomez Castro MF, McCune BT et al. TMPRSS2 and TMPRSS4 promote SARS-CoV-2 infection of human small intestinal enterocytes. Sci Immunol 2020; 5.

19 Zhang Q, Xu Y, Lee J et al. A myosin-7B-dependent endocytosis pathway mediates cellular entry of alpha-synuclein fibrils and polycation-bearing cargos. Proc Natl Acad Sci U S A 2020; 117:10865–10875.

20 Christianson HC, Belting M. Heparan sulfate proteoglycan as a cell-surface endocytosis receptor. Matrix Biol 2014; 35:51–55.

21 Cagno V, Tseligka ED, Jones ST, Tapparel C. Heparan Sulfate Proteoglycans and Viral Attachment: True receptors or adapation Bias? Viruses 2020; 11:596.

22 Milewska A, Zarebski M, Nowak P, Stozek K, Potempa J, Pyrc K. Human coronavirus NL63 utilizes heparan sulfate proteoglycans for attachment to target cells. J Virol 2014; 88:13221–13230.

23 Milewska A, Nowak P, Owczarek K et al. Entry of Human Coronavirus NL63 into the Cell. J Virol 2018; 92.

24 Lang J, Yang N, Deng J et al. Inhibition of SARS pseudovirus cell entry by lactoferrin binding to heparan sulfate proteoglycans. PLoS One 2011; 6:e23710.

25 Naskalska A, Dabrowska A, Szczepanski A, Milewska A, Jasik KP, Pyrc K. Membrane Protein of Human Coronavirus NL63 Is Responsible for Interaction with the Adhesion Receptor. J Virol 2019; 93.

26 Karpowicz RJ, Jr., Trojanowski JQ, Lee VM. Transmission of alpha-synuclein seeds in neurodegenerative disease: recent developments. Lab Invest 2019; 99:971–981.

27 Jeon S, Ko M, Lee J et al. Identification of antiviral drug candidates against SARS-CoV-2 from FDA-approved drugs. Antimicrob Agents Chemother 2020.

28 Mycroft-West C, Su D, Elli S et al. The 2019 coronavirus (SARS-CoV-2) surface protein (Spike) S1 Receptor Binding Domain undergoes conformational change upon heparin binding. bioRxiv 2020.

29 Kim SY, Jin W, Sood A et al. Characterization of heparin and severe acute respiratory syndrome-related coronavirus 2 (SARS-CoV-2) spike glycoprotein binding interactions. Antiviral Res 2020:104873.

30 Liu L, Chopra P, Li X, Wolfert MA, Tompkins SM, Boons GJ. SARS-CoV-2 spike protein binds heparan sulfate in a length- and sequence-dependent manner. bioRxiv 2020.

31 de Haan CA, Haijema BJ, Schellen P et al. Cleavage of group 1 coronavirus spike proteins: how furin cleavage is traded off against heparan sulfate binding upon cell culture adaptation. J Virol 2008; 82:6078–6083.

32 Sarrazin S, Lamanna WC, Esko JD. Heparan sulfate proteoglycans. Cold Spring Harb Perspect Biol 2011; 3.

33 Meneghetti MC, Hughes AJ, Rudd TR et al. Heparan sulfate and heparin interactions with proteins. J R Soc Interface 2015; 12:0589.

34 Song Y, Thiagarajah J, Verkman AS. Sodium and chloride concentrations, pH, and depth of airway surface liquid in distal airways. J Gen Physiol 2003; 122:511–519.

35 Park RJ, Wang T, Koundakjian D et al. A genome-wide CRISPR screen identifies a restricted set of HIV host dependency factors. Nat Genet 2017; 49:193–203.

36 Wernick NL, Chinnapen DJ, Cho JA, Lencer WI. Cholera toxin: an intracellular journey into the cytosol by way of the endoplasmic reticulum. Toxins (Basel) 2010; 2:310–325.

37 Ekins S, Madrid PB. Tilorone, a Broad-Spectrum Antiviral for Emerging Viruses. Antimicrob Agents Chemother 2020; 64.

38 Le Tourneau C, Raymond E, Faivre S. Sunitinib: a novel tyrosine kinase inhibitor. A brief review of its therapeutic potential in the treatment of renal carcinoma and gastrointestinal stromal tumors (GIST). Ther Clin Risk Manag 2007; 3:341–348.

39 Melak M, Plessner M, Grosse R. Actin visualization at a glance. J Cell Sci 2017; 130:525–530.

40 Kouznetsova J, Sun W, Martinez-Romero C et al. Identification of 53 compounds that block Ebola virus-like particle entry via a repurposing screen of approved drugs. Emerg Microbes Infect 2014; 3:e84.

41 Kapuscinski J, Darzynkiewicz Z. Interactions of antitumor agents Ametantrone and Mitoxantrone (Novatrone) with double-stranded DNA. Biochem Pharmacol 1985; 34:4203–4213.

42 Crespi MD, Ivanier SE, Genovese J, Baldi A. Mitoxantrone affects topoisomerase activities in human breast cancer cells. Biochem Biophys Res Commun 1986; 136:521–528.

43 Tandon R, Sharp JS, Zhang F et al. Effective Inhibition of SARS-CoV-2 Entry by Heparin and Enoxaparin Derivatives. bioRxiv 2020.

44 Clausen TM, Sandoval DR, Spliid CB et al. SARS-CoV-2 Infection Depends on Cellular Heparan Sulfate and ACE2. bioRxiv 2020.

45 Liu C, Tang J, Ma Y et al. Receptor usage and cell entry of porcine epidemic diarrhea coronavirus. J Virol 2015; 89:6121–6125.

46 Schwegmann-Wessels C, Zimmer G, Schroder B, Breves G, Herrler G. Binding of transmissible gastroenteritis coronavirus to brush border membrane sialoglycoproteins. J Virol 2003; 77:11846–11848.

47 Cai Y, Zhang J, Xiao T et al. Distinct conformational states of SARS-CoV-2 spike protein. Science 2020.

48 Lan J, Ge J, Yu J et al. Structure of the SARS-CoV-2 spike receptor-binding domain bound to the ACE2 receptor. Nature 2020; 581:215–220.

49 Yan R, Zhang Y, Li Y, Xia L, Guo Y, Zhou Q. Structural basis for the recognition of SARS-CoV-2 by full-length human ACE2. Science 2020; 367:1444–1448.

50 Wrapp D, Wang N, Corbett KS et al. Cryo-EM structure of the 2019-nCoV spike in the prefusion conformation. Science 2020; 367:1260–1263.

51 Henderson R, Edwards RJ, Mansouri K et al. Controlling the SARS-CoV-2 spike glycoprotein conformation. Nat Struct Mol Biol 2020.

52 Yamauchi Y, Helenius A. Virus entry at a glance. J Cell Sci 2013; 126:1289–1295.

53 Mayer GD, Krueger RF. Tilorone hydrochloride: mode of action. Science 1970; 169:1214–1215.

54 Riggs BL, Hartmann LC. Selective estrogen-receptor modulators -- mechanisms of action and application to clinical practice. N Engl J Med 2003; 348:618–629.

55 Millet JK, Tang T, Nathan L et al. Production of Pseudotyped Particles to Study Highly Pathogenic Coronaviruses in a Biosafety Level 2 Setting. J Vis Exp 2019.

56 Millet JK, Whittaker GR. Murine Leukemia Virus (MLV)-based Coronavirus Spike-pseudotyped Particle Production and Infection. Bio Protoc 2016; 6.

